# Senescence phenotype of lymph node stromal cells from patients with rheumatoid arthritis is partly restored by dasatinib treatment

**DOI:** 10.1101/2023.12.10.571042

**Authors:** T.A. de Jong, J.F. Semmelink, J.W. Bolt, C. Grasso, R.A. Hoebe, P.M. Krawczyk, L.G.M. van Baarsen

## Abstract

**Objective:** Cellular senescence is a state of proliferation arrest of cells occurring during aging. The persistence and accumulation of senescent cells has been implicated in the pathogenesis of age-related diseases like rheumatoid arthritis (RA). RA is a chronic autoimmune disease in which loss of immune tolerance and systemic autoimmunity precedes clinical onset of disease. Lymph node stromal cells (LNSCs) are important regulators of immune tolerance. Accordingly, accumulating senescent LNSCs may potentially lead to defective immune tolerance and the development of systemic autoimmune disease.

**Methods:** Human LNSCs were isolated and cultured from inguinal lymph node needle biopsies from individuals at risk of developing RA (RA-risk individuals), RA patients and seronegative healthy volunteers. Senescence hallmarks and the effect of dasatinib treatment were assessed using quantitative PCR, flow cytometry, microscopy and live-cell imaging.

**Results:** Cell size, granularity and autofluorescence were significantly higher in RA LNSCs compared with control LNSCs. Stainings indicate more senescence associated β-galactosidase activity, more lipofuscin positive granules and increased DNA damage in RA-risk and RA LNSCs compared with control LNSCs. Moreover, we found altered gene expression levels of senescence associated genes in LNSCs from RA patients. Strikingly, the capacity to repair irradiation induced DNA damage was significantly lower in RA-risk and RA LNSCs compared with control LNSCs. Treating LNSCs with dasatinib significantly improved cell size and DNA repair capacity of cultured LNSCs.

**Conclusion:** We observed multiple senescent hallmarks in RA LNSCs and to lesser extent already in RA-risk LNSCs, which could partly be restored by dasatinib treatment.

**KEY MESSAGES:** *What is already known on this topic?:*

– Synovial fibroblasts from RA patients display a senescent phenotype and accumulate in inflamed synovial tissue.

*What does this study add?:*

– Lymph node stromal cells (LNSCs) from RA patients, and to a lesser extent from RA-risk, display key hallmarks of senescence.
– Both *ex vivo* and *in vitro* LNSCs from RA patients have an increased cell size compared with control LNSCs.
– RA and RA-risk LNSCs have an impaired ability to repair DNA damage
– Treating LNSCs with dasatinib significantly improved cell size and DNA repair capacity of LNSCs.

*How might this study impact on clinical practice or future developments?:*

– These hallmarks of senescence in LNSCs may indicate premature aging and loss of function of the immunomodulatory lymph node stromal compartment during RA development. Dasatinib treatment of LNSCs shows that senolytics may be an effective preclinical drug to restore cell function early in disease.

## Introduction

Aging and age-related diseases have become more prevalent since the average human life span has drastically increased. One hallmark of aging is the accumulation of senescent cells (1). These cells are in a state in which they are viable and metabolically active yet do not proliferate. Cellular senescence can be described as an emergency brake on cells that allows for proper repair during cell cycle arrest to prevent them from tumorigenesis (2, 3). However, the persistence and accumulation of senescent cells can cause tissue imbalance and has been implicated in the pathogenesis of age-related diseases like rheumatoid arthritis (RA) (4, 5).

Aging is an important risk factor of RA, a systemic chronic autoimmune disease in which immune cells infiltrate synovial tissue (6). Age-associated changes in immune function contribute to cellular senescence and create a pro-inflammatory microenvironment which might accelerate disease progression in RA (5, 7). Previous studies have defined premature aging of T cells in RA patients and showed that maladaptive aging of T cells contributes to the development of autoimmunity (8, 9). Moreover, it has been shown that synovial fibroblasts from RA patients display pro-inflammatory and senescent characteristics and accumulate in synovial tissues compared with age-matched controls (10). These studies show that senescence in RA is not limited to one cell type. As most studies have been performed using tissues of patients with established disease, it is unclear whether senescence is a primary cause of RA or secondary to chronic inflammation.

In the absence of synovial inflammation (11, 12), RA-specific autoantibodies such as rheumatoid factor (RF) and anti-citrullinated protein antibodies (ACPAs) can be present years before clinical manifestation of disease (13–15). This allows the identification of individuals at risk of developing RA (RA-risk) (16) and enables studies on the preclinical phase of disease in absence of inflammation. Because systemic autoimmunity apparently precedes synovial tissue inflammation, it is hypothesized that other unidentified immune processes, possibly outside the synovium, are altered and contribute to disease development. Because autoimmunity can develop when tolerance mechanisms are not properly controlled in secondary lymphoid organs, this might be a location of interest to study the at-risk phase of RA. In secondary lymphoid organs, such as peripheral lymph nodes (LNs), the modulation of effective immune responses and the regulation of peripheral tolerance depend on the proper functioning of lymph node stromal cells (LNSCs) (17–20). Previously, we have shown that LN fibroblasts from patients with RA have a reduced capacity to respond to external triggers like TLR-3 and TNF (21, 22). We hypothesize that during the development of RA, LNSCs might exhibit a senescent phenotype and accumulate prematurely in lymphoid organs which can potentially lead to defective peripheral tolerance, improper control of immune responses and the development of systemic autoimmune disease.

To date, the aetiology of RA remains unclear and despite current treatment options are able to suppress inflammation and join pain, curative treatment does not exist. To target senescent cells there are several senolytic agents that selectively eliminate senescent cells by inducing apoptosis. Dasatinib, a tyrosine kinase inhibitor, has shown to attenuate arthritis symptoms in murine collagen-induced arthritis (23). Additionally, *in vitro* cultured synovial fibroblasts from RA patients showed decreased proliferation and migration capacity upon dasatinib treatment. Moreover, dasatinib treatment led to a significant increase in apoptosis of synovial fibroblasts (23). Transcriptional analysis comparing peripheral blood derived T cells from RA patients and age-matched healthy controls showed that ephrin B1 (*EFNB1*), a tyrosine kinase ligand targeted by dasatinib, was elevated in RA patients. Furthermore, *EFNB1* expression correlated with RA symptoms (24), making dasatinib an interesting drug for potential future therapies. In this study we investigated the senescence phenotype of LN fibroblasts (referred to as LNSCs) during health, systemic autoimmunity and RA. Additionally, we examined whether dasatinib treatment could restore their cellular phenotype and function.

## Materials and methods

### Study subjects and tissue samples

Individuals with arthralgia and/or a family history of RA who were positive for anti-citrullinated protein antibodies and without any evidence of arthritis upon examination were included. These individuals were considered to be at risk of developing RA (RA-risk individuals) (16, 25). For comparison, we included patients diagnosed with RA and aged-matched seronegative healthy volunteers. LN tissues were collected by ultrasound-guided inguinal LN needle core biopsy as previously described (26) after which LNSCs were isolated and expanded (22). The study was performed according to the principles of the Declaration of Helsinki (27), approved by the Institutional Review Board of the Amsterdam UMC and all study subjects gave their written informed consent. Table 1 shows the demographics of the included study subjects. Detailed description of the included study subjects and methods used for RNA extraction, quantitative real-time PCR, flow cytometry and microscopy can be found in the supplementary file.

**Table 1:** Demographic baseline characteristics of study subjects.

|  | Ex vivo cohort |  |  | Cell culture cohort |  |  |
| --- | --- | --- | --- | --- | --- | --- |
|  | Controls | RA-risk individuals | RA patients | Controls | RA-risk individuals | RA patients |
|  | n=9 | n=4 | n=11 | n=5 | n=5 | n=5 |
| Sex (female, n) (%) | 5 (56) | 2 (50) | 5 (45) | 3 (60) | 4 (80) | 4 (80) |
| Age (years) | 47 (37-63) | 51 (41-57) | 69 (59-72) | 47 (33-58) | 48 (36-50) | 42 (29-44) |
| IgM-RF positive (n) (%) | ND | 3 (75) | 7 (64) | 0 (0) | 0 (0) | 4 (80) |
| ACPA positive (n) (%) | ND | 3 (75) | 9 (82) | 0 (0) | 5 (100) | 5 (100) |
| ESR (mm/h) | ND | 6.5 (2.8-8) | 12 (5-22) | ND | 5 (2-8.5) | 26 (6.5-40.5) |
| CRP | ND | 0.7 (0.5-1.9) | 3.8 (1.1-8) | 0.5 (0.3-3.9) | 1.6 (0.8-3.6) | 3 (1.7-26.5) |

ND, not determined; IgM-RF, IgM rheumatoid factor; ACPA, anti-citrullinated protein antibodies; ESR, erythrocyte sedimentation rate; CRP, C-reactive protein; VAS, visual analogue scale. Data are expressed as median (interquartile range) unless otherwise indicated. IgM RF: IgM rheumatoid factor (measured using IgM-RF ELISA, ULN 49 IU/mL (Hycor Biomedical, Indianapolis, IN)).

## Results

### LNSCs from RA patients have an increased size and granularity

Senescent cells exhibit several morphological alterations and are described to have an enlarged and irregular cell shape (28). Flow cytometry gating strategy used to identify LNSCs is shown in Figure 1a. Quantitative analysis of LNSCs *ex vivo* revealed that RA LNSCs are significantly larger compared with control LNSCs (Figure 1b). Cell size of LNSCs from RA-risk individuals was numerically, but non-significantly, higher compared with control LNSCs. Cell granularity was not significantly different between diagnoses (Figure 1c). This data suggests that morphological alterations can be observed in LNSCs directly *ex vivo*.

**Figure 1.**
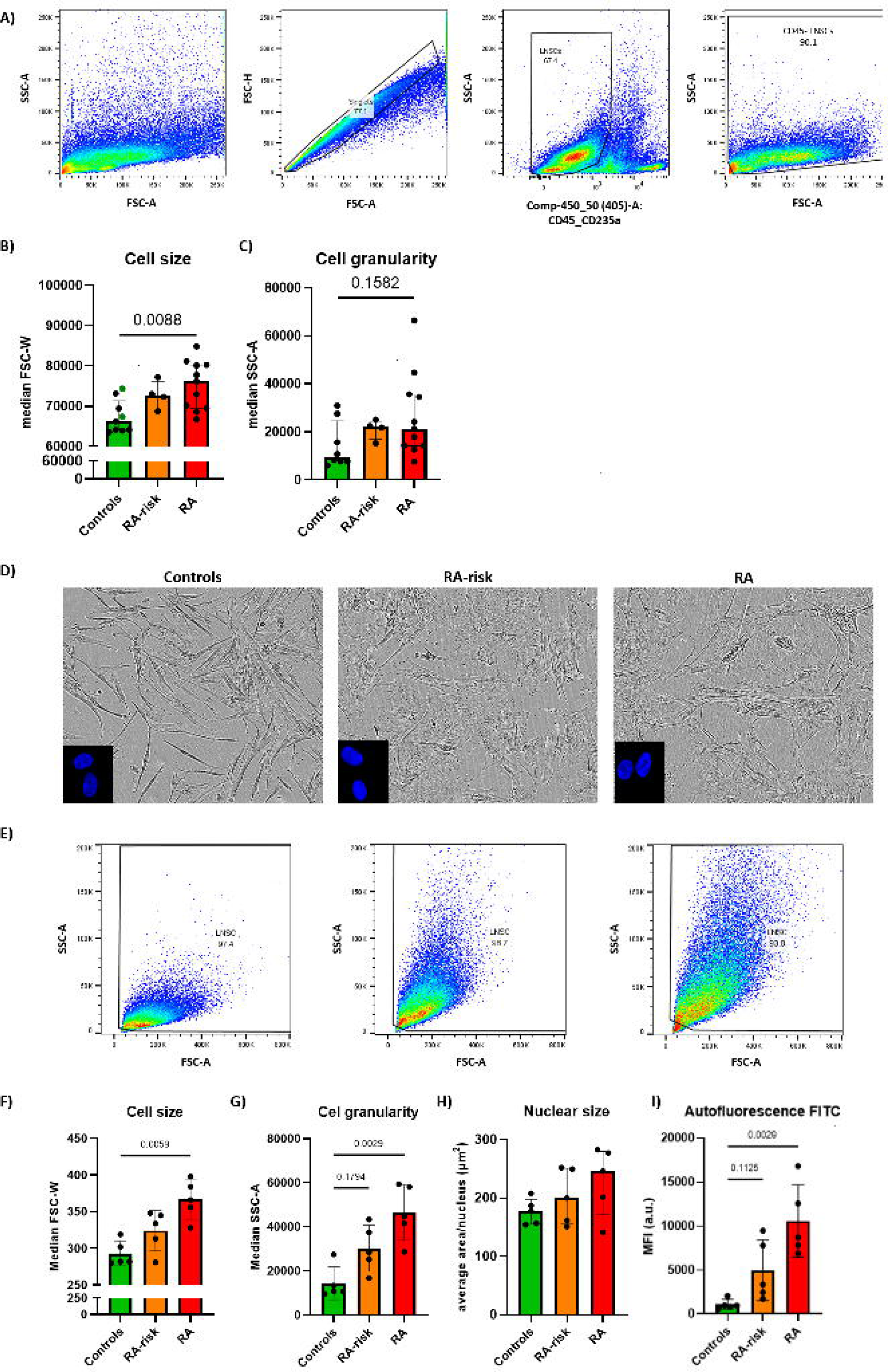
Morphological alterations in directly *ex vivo* analyzed and cultured RA LNSCs compared with control LNSCs. A) Flow cytometry gating strategy used to identify single cells and LNSCs. Numbers adjacent to the outlined areas indicate percentages of cells in the gated population. B) Cell size and C) Cell granularity measured by flow cytometry. Green = non-kidney transplant healthy volunteers undergoing LN needle biopsy. D) Representative phase-contrast images of cultured LNSCs using Incucyte ZOOM. Insert: representative immunofluorescent stainings of LNSCs nuclei using DAPI. E) Representative flow cytometry plots. F) Cell size measured by flow cytometry (MFI of FSC-W) G) Cell granularity (MFI of SSC-A). H) Nuclear size of cultured LNSCs measured by immunofluorescent DAPI staining, approximately 50 cells per donor were measured for analysis by LAS X 3D and ImageJ. I) Autofluorescence of unstained cells in FITC channel. All donors passage 5 for flow cytometry, passage 6 for nuclear size measurements. N=5 per group. Data are presented as median + interquartile range. Statistical differences were determined using a Kruskal-Wallis test followed by Dunn’s multiple comparisons test.

Additionally, *in vitro* cultured LNSCs from RA patients and RA-risk individuals also showed morphological differences when compared with control LNSCs on both microscopic images as well as spectral flow cytometry plots (Figure 1d, e). Flow cytometry measurements of unstained LNSCs show that RA LNSCs harbor a significantly larger cell size and higher granularity compared with control LNSCs (Figure 1f, g). Immunofluorescent stainings, did not show any significant differences in nuclear size between donor groups (Figure 1h). Increased autofluorescence was recently discussed as reliable senescence marker for *in vitro* cultured human mesenchymal stromal cells as it correlates with multiple established senescence markers (29). In our cohort, autofluorescence in FITC channel was significantly higher in RA LNSCs compared with control LNSCs (Figure 1i). Autofluorescence was detected at similar levels in other fluorescent channels (data not shown). Altogether, this data revealed that the *ex vivo* increase in cell size is maintained during LNSC culture allowing functional *in vitro* studies to assess additional senescence hallmarks.

### Increased lysosomal content in RA(-risk) LNSCs

We next evaluated lysosomal content of cultured LNSCs by measuring the activity senescence-associated β-galactosidase (SA-β-gal), a lysosomal enzyme. Cultured LNSCs were defined SA-β-gal positive at a cut off intensity according to the Senescence Counter macro in ImageJ as shown in figure 2a. Hardly any positive cells were detected in control LNSCs, whereas RA LNSCs displayed a relatively high percentage of blue stained cells (Figure 2b). Image quantification showed that SA-β-gal activity was significantly higher in RA LNSCs and numerically higher in RA-risk LNSCs compared with control LNSCs (Figure 2c).

**Figure 2.**
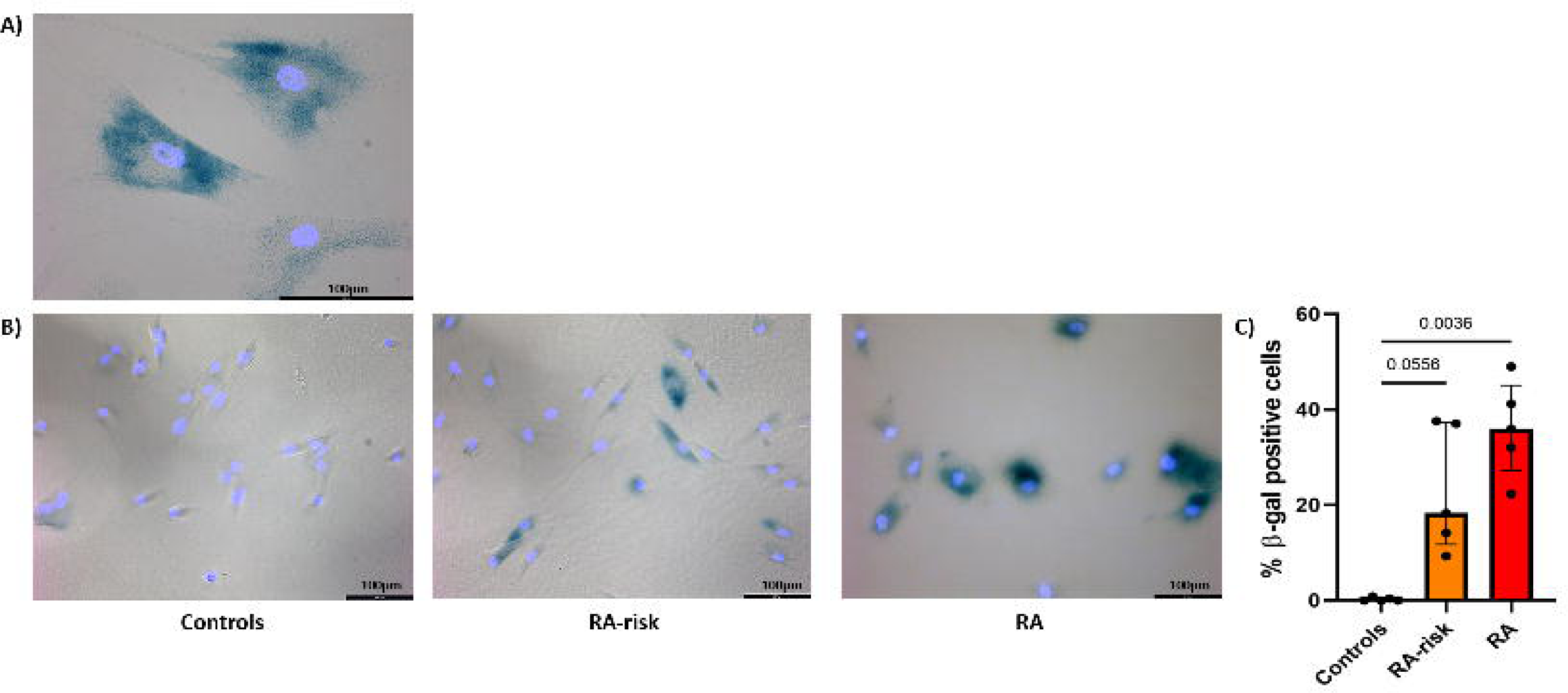
**Significantly higher SA-**β**-gal activity in RA-risk and RA LNSCs compared with control LNSCs.** A) Representative image reflecting positive and negative SA-β-gal staining when applying the Senescence Counter macro in ImageJ. B) Representative images of SA-β-gal (dark blue) combined with DAPI nuclear staining (light blue in cultured LNSCs. C) SA-β-gal positive LNSCs as % of DAPI positive cells. All donors passage 6, n=5 per group and approximately 50 cells per donor were analyzed. Data are presented as median + interquartile range. Statistical differences were determined using a Kruskal-Wallis test followed by Dunn’s multiple comparisons test.

### Morphological alterations partially restored by dasatinib treatment

Because we detected relatively high SA-β-gal positivity in cultured LNSCs, we decided to investigate whether LNSCs are sensitive to senolytics that specifically target senescent cells. The most studied senolytic in fibroblasts is Navitoclax, which induces apoptosis by targeting the anti-apoptotic pathways of BCL2 and BCL-xL. However, titration experiments in LNSCs did not show any effect on cell viability (supplementary figure 1a), suggesting that LNSCs are not sensitive for Navitoclax. In line with this, mRNA levels of *BCL2L1* (encoding for Bcl-xL) was not differentially expressed between donor groups (supplementary figure 1b). The senolytic piperlongumine decreased cell viability in both control and RA LNSCs (supplementary figure 1c), making it an unsuitable drug to target altered LNSCs in RA patients and quercetin did not show any effect (supplementary figure 1d). In contrast, dasatinib, reported to effectively remove senescent pre-adipocytes (30), did reduce cell viability only in RA LNSCs (supplementary figure 1e). Dasatinib is often used in combination with quercetin, but we did not detect an additional drop in cell viability after combining the two senolytic drugs (data not shown). Dasatinib interferes with ephrin B1 (EFNB1), an ephrin dependent receptor ligand, which was non-significantly higher expressed in RA LNSCs compared with control LNSCs (supplementary figure 1f). Guided by these findings, showing a selective effect of dasatinib treatment on RA LNSCs, we next investigated the effect of dasatinib treatment on cell morphology.

To selectively eliminate senescent cells, cultured RA-risk and RA LNSCs were treated with 5µM dasatinib for 24h. Subsequently, dasatinib was washed away after which remaining cells were given the time to proliferate until confluence of the untreated cells, followed by cell harvest and analyses (Figure 3a). To demonstrate the functional effect of dasatinib on senescent cells, healthy LNSCs were irradiated before treatment which, as expected, led to a significant increase in cell and nuclear size and a numerical increase in cell granularity and autofluorescence (Figures 3b, c, d, e and supplementary figure 2). Dasatinib treatment partly restored the morphological alterations observed in RA-risk and RA patients, as well as in irradiated healthy LNSCs (Figures 3b, c, d, and e).

**Figure 3.**
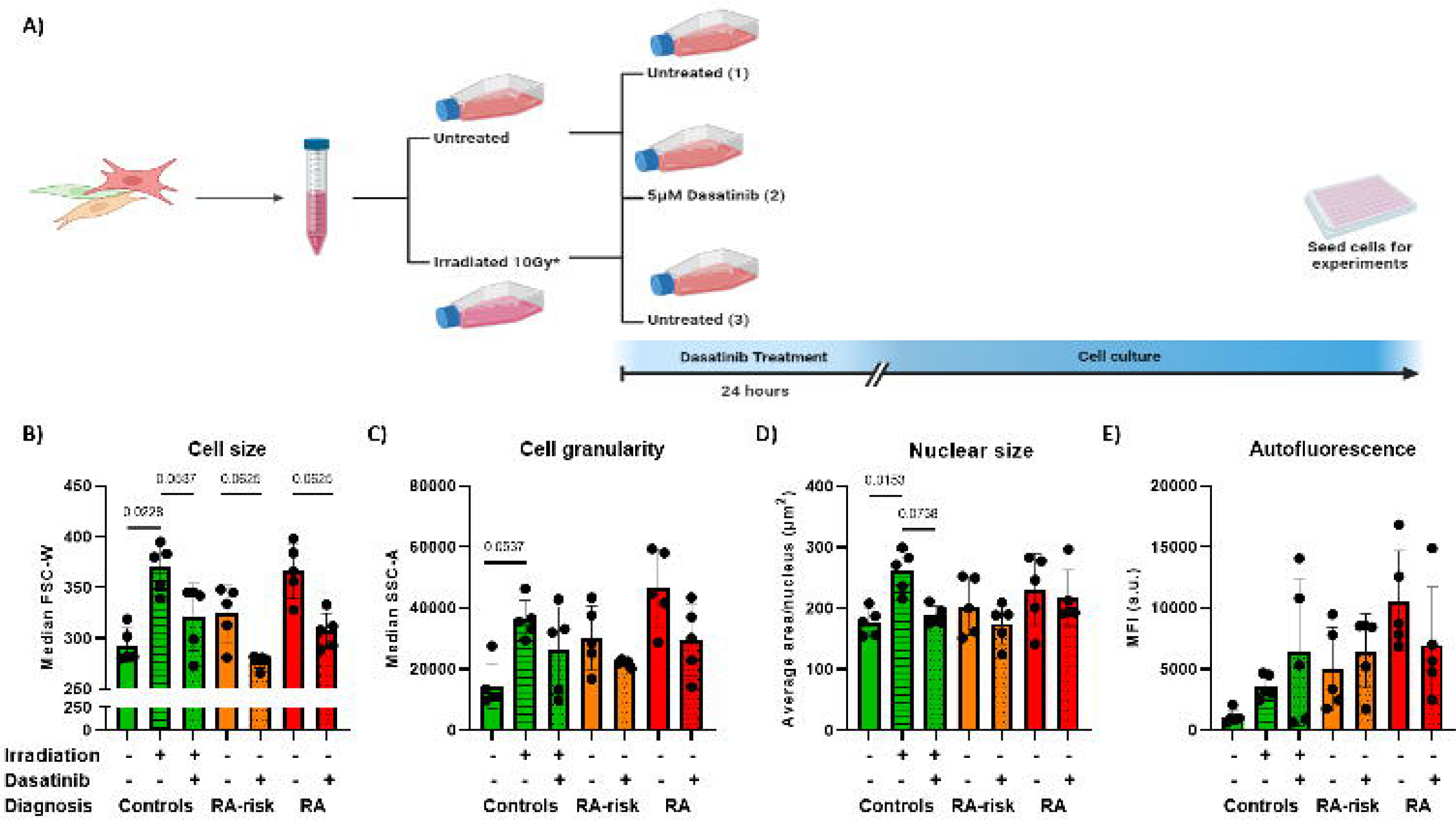
Dasatinib treatment partially restores cell size in RA-risk and RA LNSCs. A) Graphical overview of dasatinib treatment experimental setup. After cell collection, RA-risk and RA LNSCs were seeded in two flasks and control LNSCs in three flasks. The next day, one flask was treated with 5µM dasatinib for 24h and the other was left untreated. After 24h, dasatinib was washed away and remaining cells were left in culture until flasks with untreated LNSCs reached 80% confluence. Upon reaching confluence, both flasks per donor were harvested and seeded for experiments. Two flasks of control LNSCs were irradiated with 10 gray, from which one was treated for 24 hours with 5µM dasatinib 24 hours after irradiation and the other was left only irradiated. The third control LNSCs flask was untreated and once these LNSCs were confluent, the other two flasks were also harvested and plated out for experiments. *Only control LNSCs. B) Cell size C) Cell granularity D) Nuclear size (approximately 50 cells per donor were measured for analysis) and E) autofluorescence of RA-risk and RA LNSCs before and after dasatinib treatment and control LNSCs before and after irradiation and dasatinib treatment. All donors passage 5 for flow cytometry, passage 6 for nuclear size measurements. Data are presented as median + interquartile range and a Wilcoxon matched pairs signed rank test was used to analyze the effect of dasatinib treatment.

### Dasatinib treatment restores increased lysosomal accumulation in LNSCs

We next evaluated whether the observed increased SA-β-gal activity in RA(-risk) LNSCs would also decrease after dasatinib treatment. A significant increase of SA-β-gal positive cells in irradiated control LNSCs confirms that those cells have adapted a senescence phenotype (Figure 4a). Dasatinib treatment after irradiation significantly reduced the percentage of SA-β-gal positive cells in these control LNSCs. Although dasatinib treatment did not show a significant effect on SA-β-gal activity in both RA and RA-risk LNSCs, there was a pronounced decline in the expression levels of *GLB1*, the gene coding for SA-β-gal (Figure 4b).

**Figure 4.**
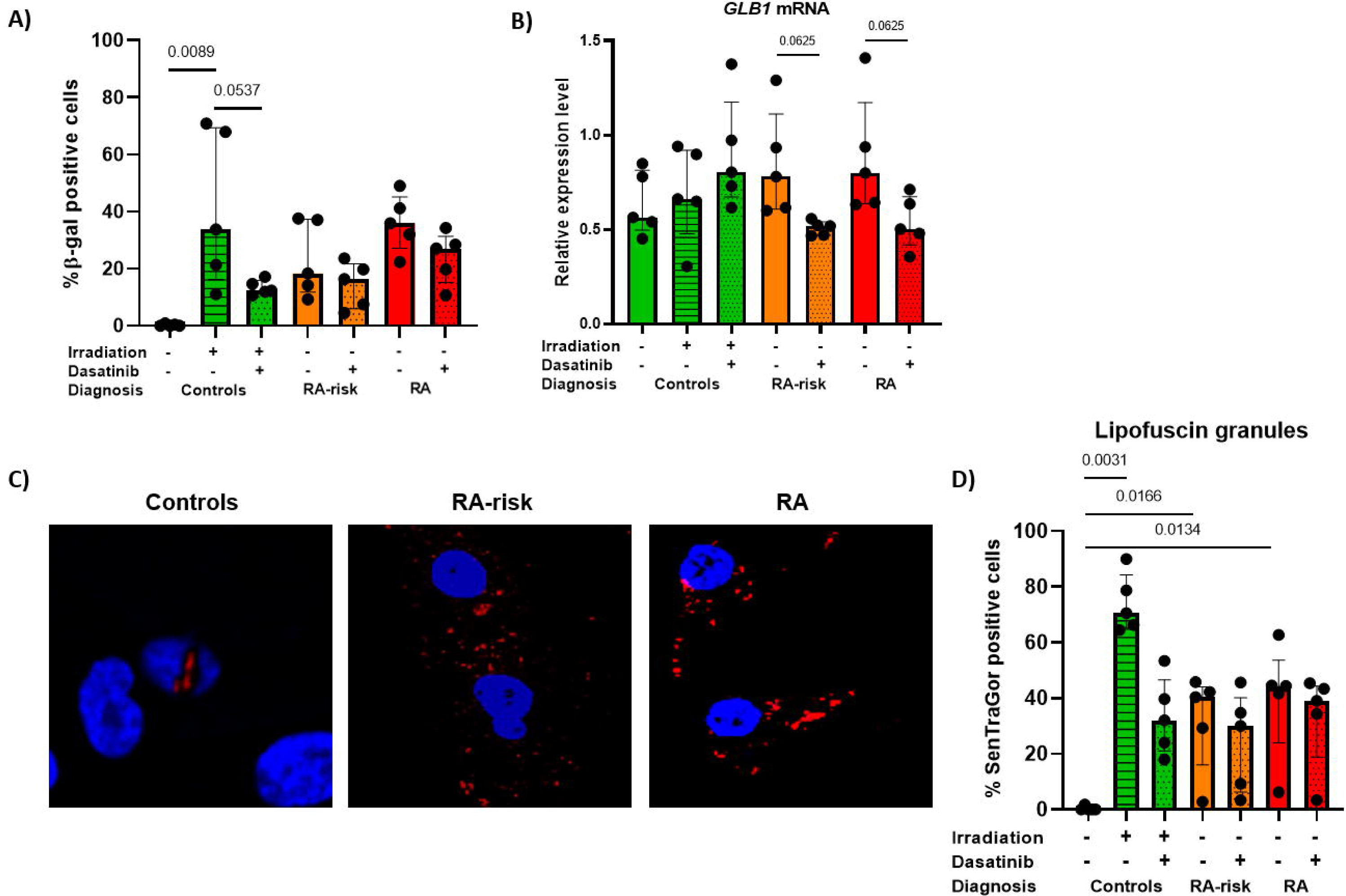
Dasatinib treatment restores increased lysosomal accumulation in LNSCs. A) SA-β-gal positive LNSCs as % of DAPI positive cells before and after treatment with 5µM dasatinib. B) Relative gene expression level of *GLB1* C) Representative images of SenTraGor staining lipofuscin granules (red) and DAPI staining nuclei (blue) in cultured LNSCs. D) Percentage of SenTraGor positive cells relative to DAPI positive cells as measured by LAS X 3D and ImageJ. All donors passage 6, N=5 per group, approximately 50 cells per condition were analyzed. Data are presented as median + interquartile range. Statistical differences at baseline were determined using a Kruskal-Wallis test followed by Dunn’s multiple comparisons test and a Wilcoxon matched pairs signed rank test was used to analyze the effect of dasatinib treatment.

Another marker for lysosomal biogenesis is SenTraGor, used to detect lipofuscin granules. Lipofuscin granule accumulation indicates reduced lysosomal degradation. SenTraGor can be visualized using immunofluorescence and was hardly detected in control LNSCs (Figure 4c). In line with the SA-β-gal data, irradiation of control LNSCs significantly increased the amount of lipofuscin granules which was reduced after dasatinib treatment (Figure 4d), suggesting that dasatinib treatment is able to target lysosomes. The percentage of SenTraGor positive cells was significantly higher in both RA-risk and RA LNSCs compared with control LNSCs. The effectiveness of dasatinib to target endogenous lipofuscin granules was highly variable within donor groups (Figure 4d).

### Senescence-associated gene expression profile in RA LNSCs

Senescent cells accumulate primarily in tissues with chronic inflammation which can have detrimental consequences for tissue structure and regeneration (31, 32). Senescent cells are able to create a pro-inflammatory microenvironment by secreting a variety of different molecules to communicate with adjacent cells, such as interleukins, chemokines, growth factors and growth regulators. Released factors can affect cell proliferation, disruption of tissue structure and function and immunomodulation (33, 34). Indeed, gene expression levels of TP53, CDKN1A, IL6, FOXO4 and CD38 are significantly higher expressed in RA LNSCs compared with control LNSCs (Figure 5). CD38 is related to coenzyme NAD, a critical metabolic for many cellular processes, other genes linked to NAD, *SIRT1, PARP1* and *NAMPT* were not differentially expressed between donor groups (supplementary figure 3). Expression of *NOTCH3* was already significantly altered in RA-risk LNSCs (Figure 5). Dasatinib treatment had no significant effect on gene expression levels in cultured LNSCs, although dasatinib did restore expression levels of *TP53*, *CDKN2A, CDKN1A* and *FOXO4* in RA(-risk) LNSCs to the expression levels measured in control LNSCs and partially restored the expression levels of *IL6*, *NOTCH3*, *LMNB1* and *CD38*. This shows that also at transcriptional level senescence can be detected in cultured LNSCs of RA patients, which is restored after dasatinib treatment for most of the analyzed genes.

**Figure 5.**
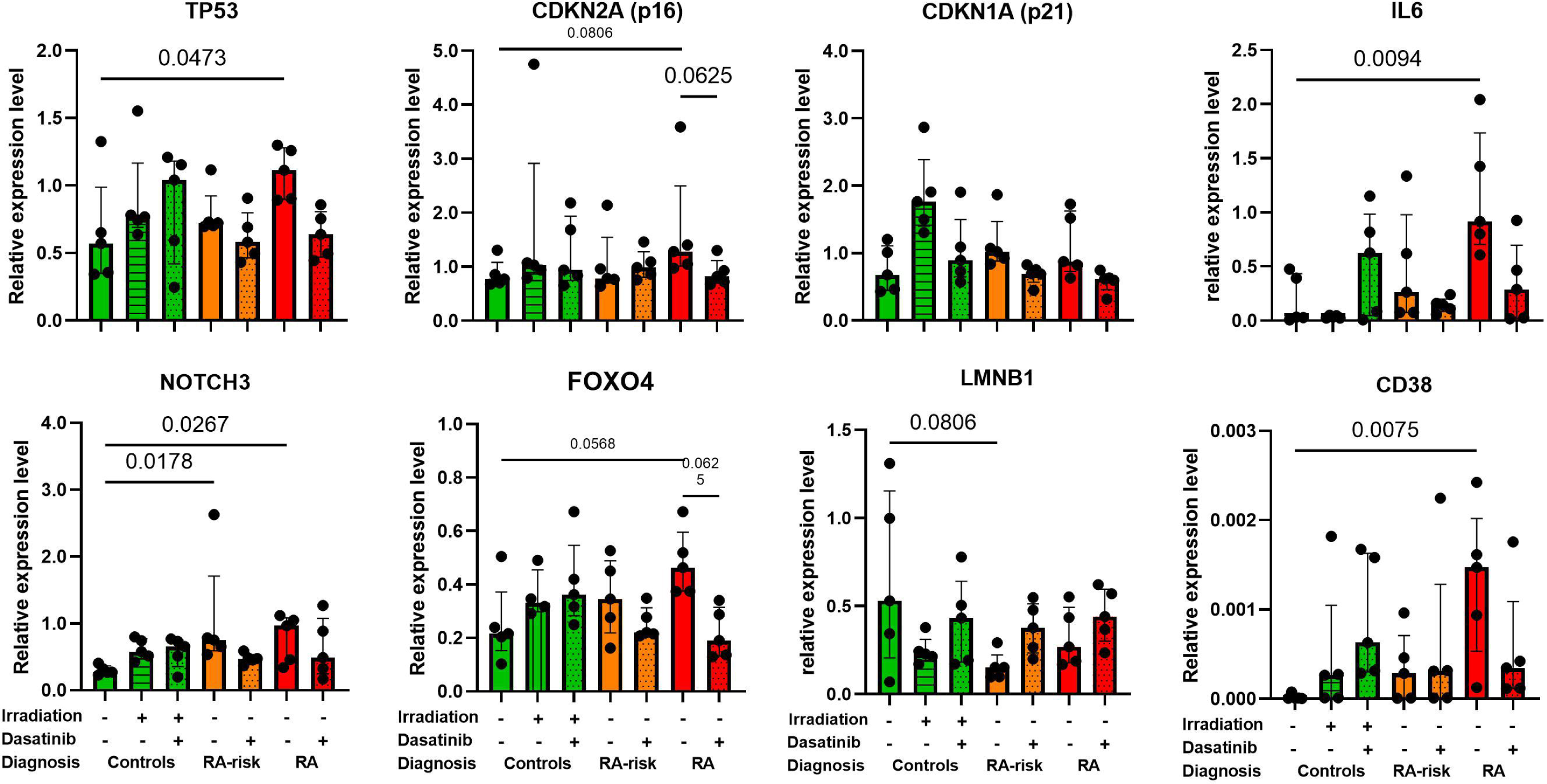
Altered gene expression profiles in RA(-risk) LNSCs. Gene expression levels of senescence associated genes in cultured LNSCs. All donors passage 6, N=5 per group. Data are presented as median + interquartile range. Statistical differences at baseline were determined using a Kruskal-Wallis test followed by Dunn’s multiple comparisons test and a Wilcoxon matched pairs signed rank test was used to analyze the effect of dasatinib treatment.

### Lower migration capacity in RA LNSCs

Next we investigated the viability, proliferation and migration of cultured LNSCs over time. Previously we have shown that the first 27 hours after seeding, LNSCs are only attaching and spreading, while the proliferation phase starts after these 27 hours (35). As the viability of the cells is similar between controls and RA(-risk) LNSCs 24 hours after seeding, (Figure 6a), we conclude that, as expected, a similar amount of LNSCs has been seeded for all donors tested. Over time, viability of all groups increase, indicating cell proliferation.

**Figure 6.**
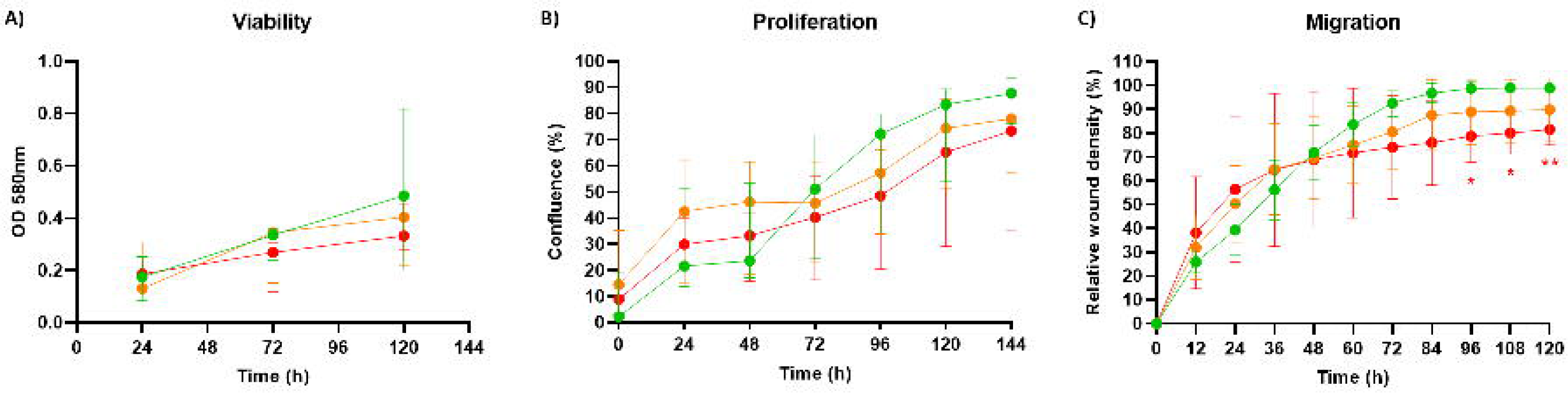
RA LNSCs have a lower migration capacity compared with control LNSCs. A) Viability of cultured LNSCs measured using MTT. B) Real-time cellular proliferation of cultured LNSCs measured using IncuCyte software. C) Real-time cellular migration capacity of cultured LNSCs measured using scratch wound assay software from IncuCyte. All donors passage 6, N=5 per group, 3 technical replicates per condition for viability and 5 technical replicates for proliferation and migration assays. Data are presented as median + interquartile range. Statistical differences were determined using a repeated measures ANOVA with the Geisser-Greenhouse correction followed by Dunnett’s multiple comparisons test. Statistical significance is indicated as * P<0.05 and ** P<0.01.

To get more insight into the proliferation rate of LNSCs in culture, we followed the cells real-time using an IncuCyte ZOOM S3. This system automatically acquired an image every 24 hours during a 6-days cell culture experiment and analyses cell confluency as a measure of proliferation rate (Figure 6b). This life-cell analysis confirmed the slow proliferation of LNSCs, which started in control LNSCs 48h after seeding. Overall, there was no statistically significant difference in proliferation rate, but RA LNSCs tend to lag behind. As a measure of cell function, we also studied the migration capacity of cultured LNSCs using a scratch wound assay. Cells were seeded at confluence and after 24h, an equal scratch wound was made automatically in every well using the IncuCyte wound maker. To analyse the migration capacity, the relative wound density was measured every 12 hours for 5 days, representing the closing of the wound. Control LNSCs were able to close the wound after 3 days culture, while RA LNSCs never closed the wound reflecting a significantly lower migration (and/or proliferation) capacity compared with control LNSCs (Figure 6c).

We next investigated whether dasatinib could restore these observed alterations in cell behavior. As expected, irradiation of control LNSCs had a negative impact on their viability, migration and proliferation and dasatinib treatment could restore part of these defects (supplementary figure 4). However, dasatinib treatment in RA(-risk) LNSCs did not impact their viability, proliferation or migration (supplementary figure 4). In summary, these live-cell imaging based analysis revealed that RA LNSCs have a lower migration and proliferation capacity compared with LNSCs which could not be restored by dasatinib treatment.

### RA-risk and RA LNSCs have an impaired capacity to repair DNA damage

We next quantified the number of yH2AX foci, which accumulate at sites of unresolved DNA damage. A significantly higher number of foci was observed in nuclei of RA LNSCs compared with control LNSCs, as well as a non-significant (P=0,0665) numerical increase in RA-risk LNSCs (Figure 7a, b). As positive control we irradiated LNSCs from controls, which significantly increased the number of yH2AX foci. Treating these irradiated control LNSCs for 24h with 5µM dasatinib showed a significant reduction in yH2AX foci, although not all DNA damage was repaired. This indicates that dasatinib treatment can partially restore irradiation induced DNA damage in cultured LNSCs. However, dasatinib treatment did not decrease DNA damage present in RA-risk and RA LNSCs (Figure 7b). Because we already detected DNA damage in cultured LNSCs at baseline, we next investigated whether RA(-risk) LNSCs had an impaired DNA repair capacity. Therefore, cultured LNSCs were first irradiated with 1 Gray after which DNA damage repair was followed over time by imaging these cells directly (0h), 20 and 40 hours after irradiation. Irradiation induced significantly more DNA damage in RA LNSCs compared with control LNSCs (Figure 7c, d), indicating that RA LNSCs might be more sensitive to DNA damage induction. After 20 hours DNA damage was significantly reduced in all donors, although yH2AX foci levels remained significantly higher in RA-risk and RA LNSCs compared with control LNSCs (Figure 7c, d), indicating that DNA repair is faster in control LNSCs. Complete DNA damage repair was observed in control LNSCs 40 hours after irradiation, while yH2AX foci persist in RA-risk and RA LNSCs. The number of yH2AX foci detected 40 hours after irradiation of RA-risk and RA LNSCs was still higher than measured in untreated LNSCs, indicating that DNA damage repair in these cells is slow and incomplete. Pre-treatment of cultured LNSCs with 5µM dasatinib showed significantly less yH2AX foci directly after irradiation, implying that this makes RA-risk and RA LNSCs less sensitive to irradiation induced DNA damage (supplementary figure 5). Furthermore, less yH2AX foci were detected in dasatinib pre-treated LNSCs 20 hours after irradiation and after 40 hours DNA damage was completely restored. This indicates that dasatinib pre-treatment can increase repair of irradiation induced DNA damage.

**Figure 7.**
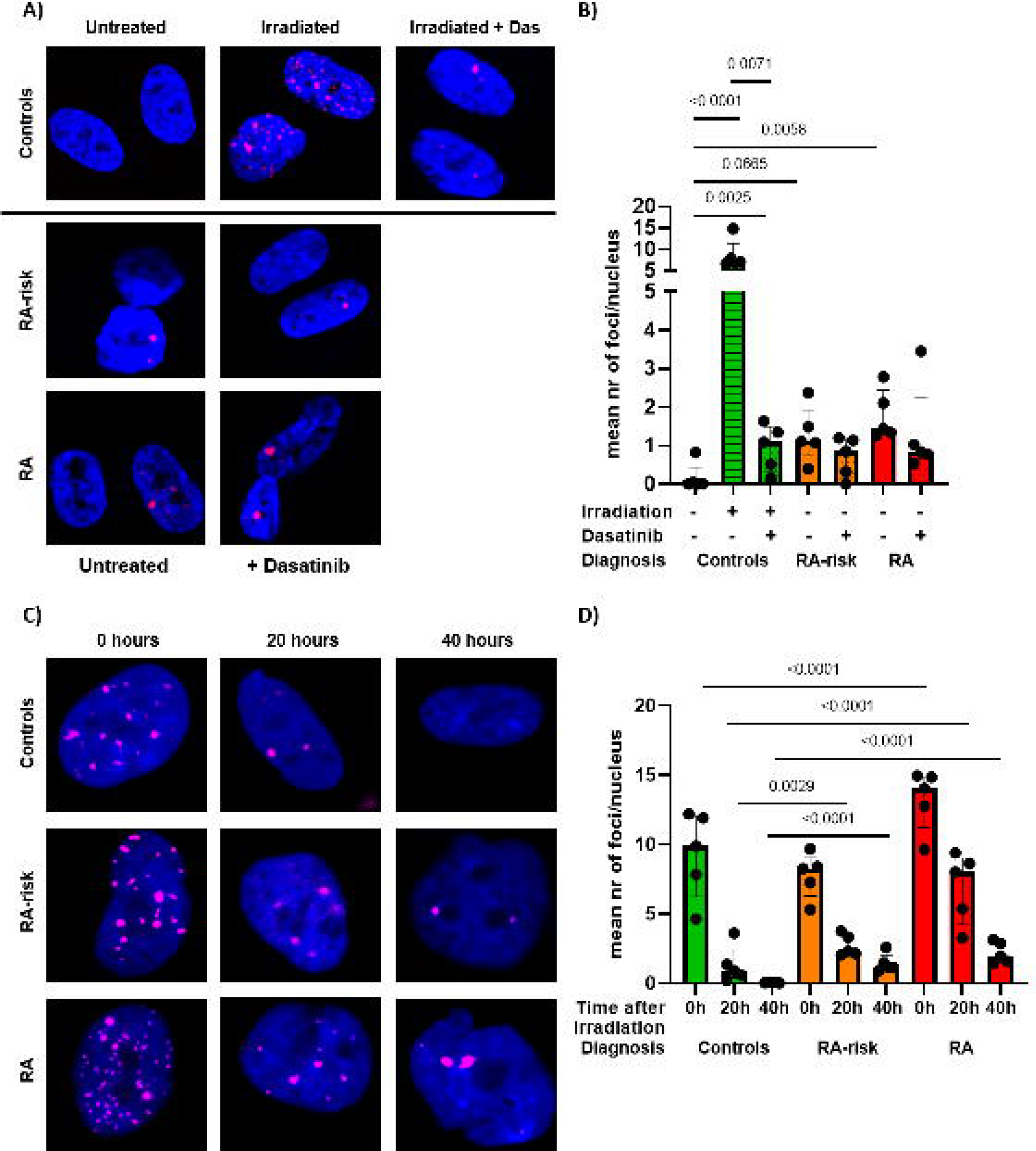
Impaired DNA damage repair in RA(-risk) LNSCs. A) Representative images of yH2AX foci (pink) and DAPI staining (blue) in cultured LNSCs. B) Average number of yH2AX foci per nucleus in cultured LNSCs before and after 5µM dasatinib treatment. All donors passage 6, unstimulated conditions. C) Representative images of yH2AX foci (pink) and DAPI staining (blue) in cultured LNSCs after DNA damage induction by gamma-irradiation. B) Average number of yH2AX foci per nucleus in cultured LNSCs directly, 20 hours and 40 hours after irradiation. Mean value per donor was determined through quantification of Z-stack images of approximately 50 cells per donor. All donors passage 7, N=5 per group. Data are presented as median + interquartile range. Statistical differences were determined using 2-way ANOVA + Dunnett’s T3 multiple comparisons test and a Wilcoxon matched pairs signed rank test was used to analyze the effect of dasatinib treatment.

## Discussion

Previous studies have shown premature cellular aging in RA. Changes detected in RA patients include accelerated immunosenescence, decreased thymic functionality, telomeric attrition and excessive senescence-associated secretory phenotype related cytokine production (5, 9, 36). Besides many T cell related studies (8, 37), it was recently shown that senescent synovial fibroblasts prematurely accumulate in synovial tissue of RA patients (10). Currently, it is unclear whether this premature aging is a primary event occurring already before onset of disease or whether it is a secondary event driven by inflammatory processes or treatment. Moreover, it is unknown whether premature aging originates in lymphoid organs crucial for immune cell differentiation and development and whether it can be observed in tissue-resident lymph node fibroblasts, which have a major influence on immune cell function.

In this study we aimed to delineate the potential senescence phenotype of LN fibroblasts during the earliest phases of autoimmunity. To investigate hallmarks of senescence during systemic autoimmunity in absence of chronic inflammation we compared LNSCs collected from autoantibody positive individuals at risk of developing RA with LNSCs from RA patients and seronegative healthy volunteers. We detected senescence markers in LNSCs from RA patients at morphological, transcriptional and functional level compared with LNSCs from age-matched controls. LNSCs from RA-risk individuals also presented several senescence markers although to a lesser extent, suggesting that senescent cell accumulation starts before onset of clinical disease

Morphological characteristics related to senescence, such as enlarged and more granular cells, were significantly more present in *ex vivo* analyzed RA LNSCs and this morphology was maintained in culture allowing functional *in vitro* studies. Moreover, cell size and granularity of cultured RA-risk LNSCs were also higher compared with control LNSCs, although to a lesser extent. Forward-sideward scatter plots suggest that there might be a smaller number of larger (senescent) cells present in RA-risk individuals compared with RA LNSCs, suggesting that accumulation of larger cells starts before onset of RA and continues after diagnosis. Increased endogenous autofluorescence in senescent and slow dividing cells has been attributed to lipofuscin accumulation in lysosomes as result of oxidative stress and ROS production (29, 38, 39). These lipofuscin granules contain highly oxidized proteins, mostly unsaturated fatty acids, that cannot be removed by proteasomal degradation and accumulate in lysosomes with age (40, 41). We previously showed increased ROS production by RA LNSCs (42) and here we observed significantly more lipofuscin granules already in RA-risk LNSCs. In aggregate, this indicates that detected autofluorescence in RA(– risk) LNSCs may reflect increased lysosomal content and oxidative stress.

In senescent cells increased lysosomal mass is partially reflected by elevated SA-β-gal activity in senescent cells (41, 43). SA-β-gal activity has been acknowledged as one of the first markers of senescence but has been extensively debated. Despite the high percentage of positivity in cultured RA LNSCs, this is not necessarily representative for *in vivo* LN tissues. Most studies comparing *in vivo* and *ex vivo* SA-β-gal stainings, found a much lower percentage of positive cells in tissue, indicating that the culture process may influence SA-β-gal activity. However, our findings can still be relevant as our age-matched control LNSCs almost don’t exhibit SA-β-gal positive cells. Moreover, our SenTraGor immunofluorescence staining was significantly higher already in RA-risk LNSCs, pointing towards accumulation of lysosomal content.

Transcriptomic analysis of senescence-associated genes in cultured LNSCs revealed multiple differentially expressed genes in RA LNSCs compared with control LNSCs. Overexpression of *FOXO4* mRNA was detected in RA LNSCs and is suggested to prevent cell death of senescent cells by sequestering p53 in the nucleus. Of note, increased *NOTCH3* expression was already detected in RA-risk LNSCs. Studies in inflamed synovium has shown that Notch3 is involved in the differentiation and arthrogenic expansion of a specific CD90+ fibroblast subset (44). Another gene upregulated in RA LNSCs is *CD38*. CD38 levels inversely correlate with the age-related decline in nicotinamide adenine dinucleotide (NAD), which ultimately leads to mitochondrial dysfunction and metabolic alterations (45, 46). It is interesting to postulate that our earlier detected metabolic alteration in LNSCs (42) is related to altered CD38 and NAD+ levels. Additional research is needed to investigate whether the differential expression of these genes are causative related to the observed senescence phenotype of RA LNSCs and may guide future studies aimed at restoring LNSC function.

Cellular senescence has been described as a state of proliferation arrest of cells. Although LNSCs proliferate slowly, we did not detect a proliferation arrest or significant differences between donor groups. However, the expression level of cyclin-dependent kinase inhibitor *CDKN2A* was significantly increased in RA LNSCs compared with control LNSCs. CDKN2A is one of the main drivers of cell cycle arrest in senescent cells (28). The increased cell size measured using flow cytometry might explain why we were not able to detect any differences in cell proliferation as this was measured using the IncuCyte, which uses confluence as proliferation readout. When cells are larger, confluence is reached earlier compared with smaller cells thus cell size is a confounding factor in our comparison between control and RA LNSCs. The scratch wound assay showed that RA LNSCs have a decreased migration capacity reflected by impaired wound repair, though this might also be caused by a difference in proliferation. Further research is needed to clarify whether or not there is a difference in proliferation rate between RA and control LNSCs.

Accumulating DNA damage or dysfunctional DNA repair machinery is linked to senescence and has been observed in T cells of patients with RA (37, 47–49). T cells closely interact with resident stromal cells within LN and synovial tissue, an interaction that might play an important role in initiating chronic inflammation. DNA damage is already present in LNSCs of RA-risk LNSCs, thus before onset of disease. The persistence of DNA damage in senescent cells activates phosphorylation of transcription factor p53, which induces growth arrest to allow for DNA damage repair. Indeed, p53 expression was significantly upregulated in RA LNSCs compared with control LNSCs. Additionally, DNA damage repair capacity was diminished in RA-risk and RA LNSCs. Despite RA-risk and control LNSCs having a similar number of yH2AX foci directly after irradiation, yH2AX foci persisted in RA-risk LNSCs after irradiation. Further research should be undertaken to investigate whether impaired DNA damage repair correlates with future arthritis development in RA-risk individuals and what effect DNA damage in LNSCs has on peripheral tolerance.

After testing four different senolytic agents, only dasatinib was able to selectively remove (senescent) LNSCs in RA patients and not affecting control LNSCs. When evaluating cell viability upon dasatinib treatment we showed that viability of irradiated control LNSCs increased after 48-72h, suggesting that irradiation-induced senescent cells were eliminated. The exponential curve after this time point suggests that healthy LNSCs then got the opportunity to grow. This effect was less clear in dasatinib treated RA LNSCs. However, dasatinib treatment did partly restore cell size and gene expression levels in RA(-risk) LNSCs. Dasatinib only had minor effect on the lysosomal content of cultured LNSCs. Data from baseline and irradiation experiments showed that dasatinib significantly improved irradiation-induced DNA damage but not the endogenous DNA damage observed in untreated RA(-risk) LNSCs suggesting different repair mechanisms are involved. The mechanism by which dasatinib (partly) restores altered senescence associated characteristics in LNSCs remains to be further elucidated. Nevertheless, dasatinib treatment of LNSCs has revealed that senolytic agents can be effective to restore LNSC function early in disease.

To our knowledge, this is the first study investigating cellular senescence in human LNSCs during systemic autoimmunity. The presence of increased numbers of senescent LNSCs in RA patients compared with controls suggest that besides T cells and synovial fibroblasts, also LN fibroblasts present signs of premature aging. Unfortunately, the relatively low number of donors tested in this study makes it hard to reach statistical significance. However, these unique human lymph node biopsy studies are extremely challenging to perform, especially in the context of systemic autoimmunity. Moreover, it is difficult to find seronegative healthy volunteers who would like to participate in such a study and LNSCs are typically slowly growing cells making these experiments very labor intensive and time consuming. A limitation of our study is that one type of senescence known as replicative senescence can easily be induced in *in vitro* culture systems. Replicative senescence is characterized by a decrease in proliferation potential induced in cells after undergoing multiple cell divisions, leading to replicative exhaustion. However, we cautiously monitored our proliferation and passaging process and compared LNSCs from different donors always from the same passage after *ex vivo* digestion. Of importance, our data in LN biopsies confirm increased cell size of RA fibroblasts analyzed directly *ex vivo* before *in vitro* expansion. Accordingly, this suggest that the observed senescence phenotype in RA(-risk) LNSCs is not purely induced by *in vitro* culture.

In conclusion, our results showed that RA LNSCs exhibit many characteristics linked to cellular senescence which could be partly restored by dasatinib treatment. Although one of the most important hallmarks of senescence, proliferation arrest, was not detected we postulate that LNSCs from RA patients display a senescence phenotype which is even partly present before onset of disease and contributes to premature aging of the LN stromal microenvironment. Further studies are needed to investigate how senescence of the local stromal microenvironment affects immune responses in the context of autoimmunity.

## Figure legends

**Supplementary figure 1.**
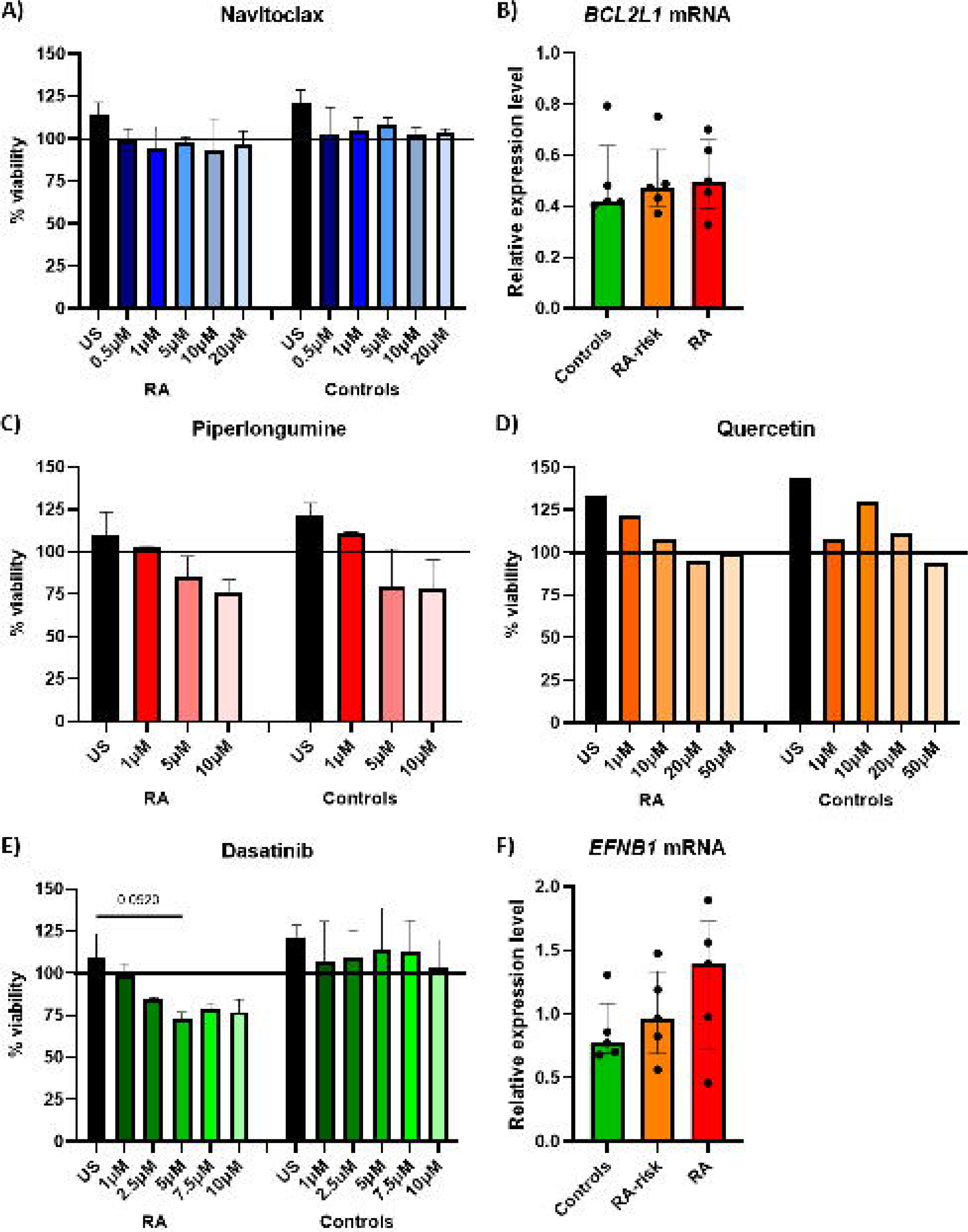
Selecting potential senolytics for LNSCs. A) Viability of RA and control LNSCs after navitoclax treatment for 24 hours. B) Relative expression level of *BCL2L1* mRNA. A gene related to the anti-apoptotic pathway of BCL-XL and a target of navitoclax. C) Viability of LNSCs after piperlongumine treatment for 24h. D) Viability of LNSCs after quercetin treatment for 24h. E) Viability of LNSCs after dasatinib treatment for 24h. Viability was measured using MTT assays. F) Relative expression level of *EFNB1* mRNA, a gene encoding for ephrin B1 which is a type I membrane protein and ligand for dasatinib. All donors passage 6 for qPCR analysis, N=5 per group. Data are presented as median + interquartile range. Statistical differences were determined using a Kruskal-Wallis followed by Dunn’s multiple comparisons test.

**Supplementary figure 2.**
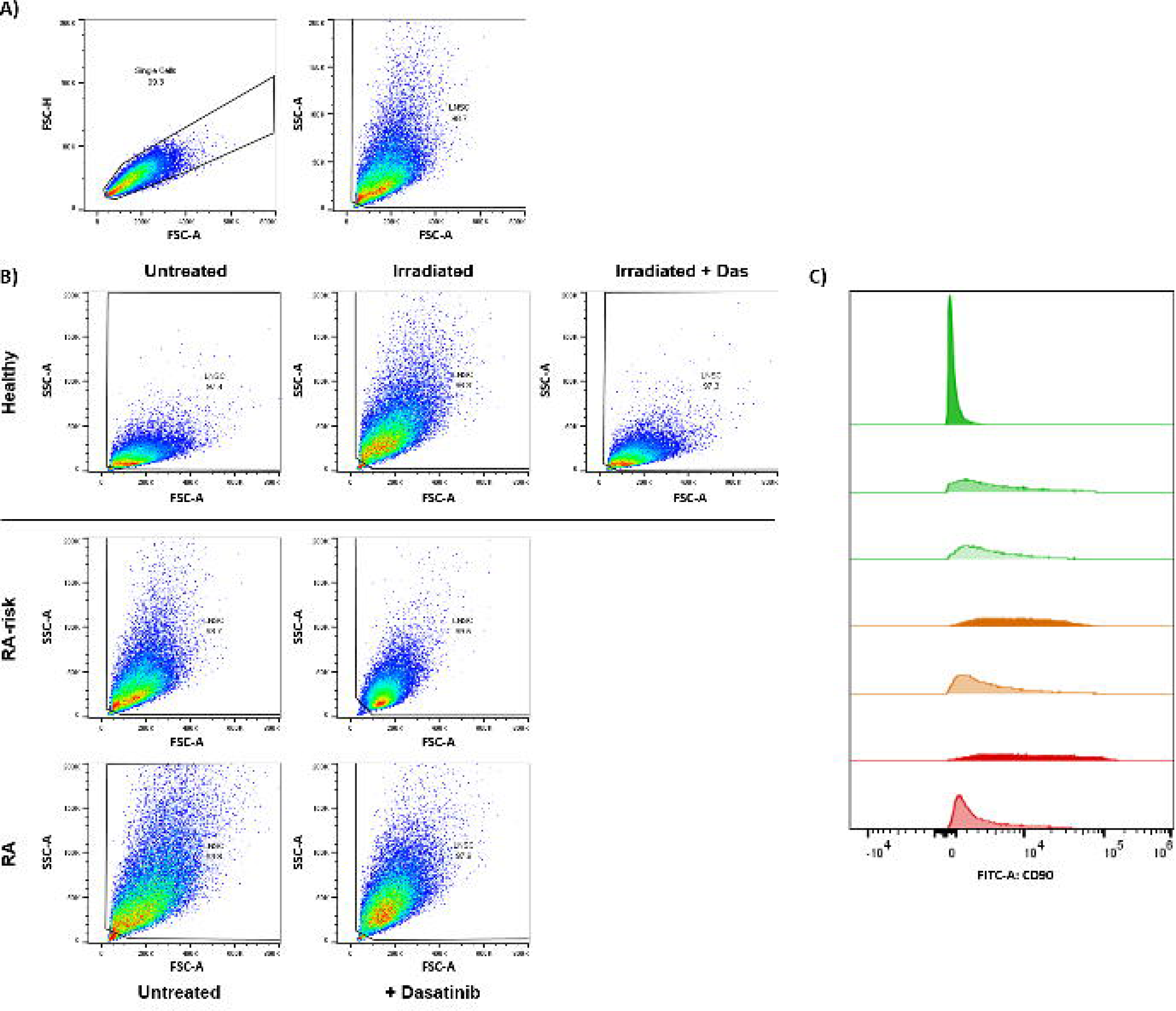
Gating strategy and flow cytometry plots of representative donors. A) Flow cytometry gating strategy used to identify single cells and LNSCs. Numbers adjacent to the outlined areas indicate percentages of cells in the gated population. B) Representative flow cytometry plots of irradiated and dasatinib (das) treated LNSCs. C) Representative mean fluorescent intensity (MFI) plots of FITC autofluorescence in unstained LNSCs. Controls (green), RA-risk individuals (orange) and RA patients (red).

**Supplementary figure 3.**
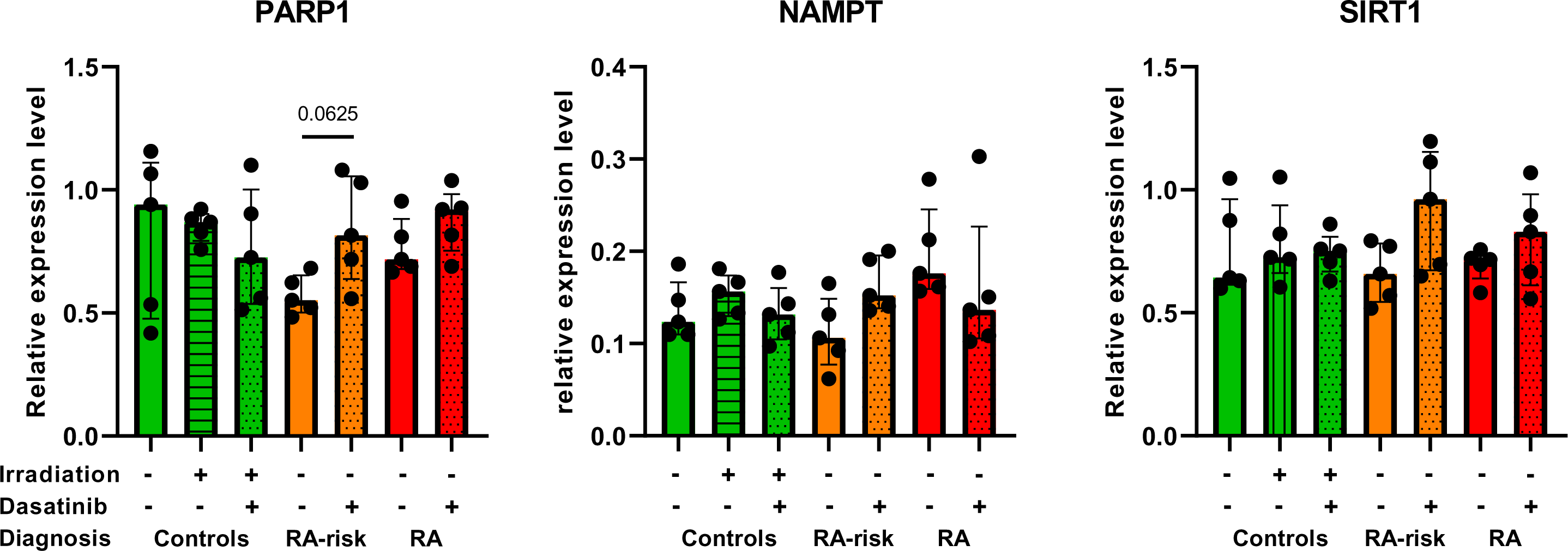
Gene expression profiles in LNSCs. Gene expression levels of senescence associated genes in cultured LNSCs. All donors passage 6, N=5 per group. Data are presented as median + interquartile range. Statistical differences at baseline were determined using a Kruskal-Wallis test followed by Dunn’s multiple comparisons test and a Wilcoxon matched pairs signed rank test was used to analyze the effect of dasatinib treatment.

**Supplementary figure 4.**
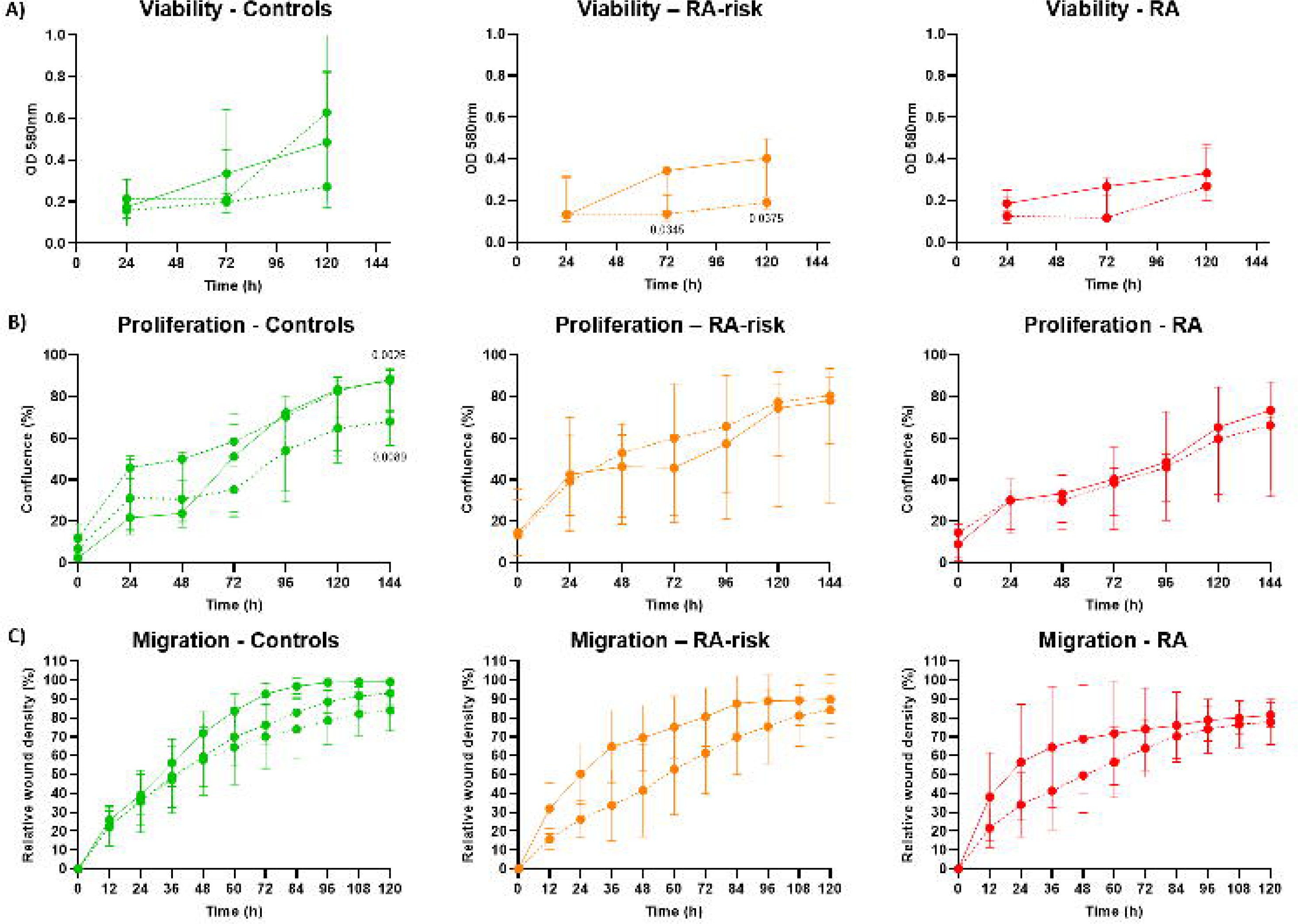
Dasatinib treatment significantly reduces cell viability in RA-risk LNSCs. A) Viability of irradiated and dasatinib treated LNSCs measured using MTT. B) Real-time cellular proliferation of irradiated and dasatinib treated LNSCs measured using IncuCyte software. C) Real-time cellular migration capacity of irradiated and dasatinib treated LNSCs measured using scratch wound assay software from IncuCyte. All donors passage 6, N=5 per group, 3 technical replicates per condition for viability and 5 technical replicates for proliferation and migration assays. Data are presented as median + interquartile range. Statistical differences were determined using a repeated measures ANOVA with the Geisser-Greenhouse correction followed by Dunnett’s multiple comparisons test.

**Supplementary figure 5.**
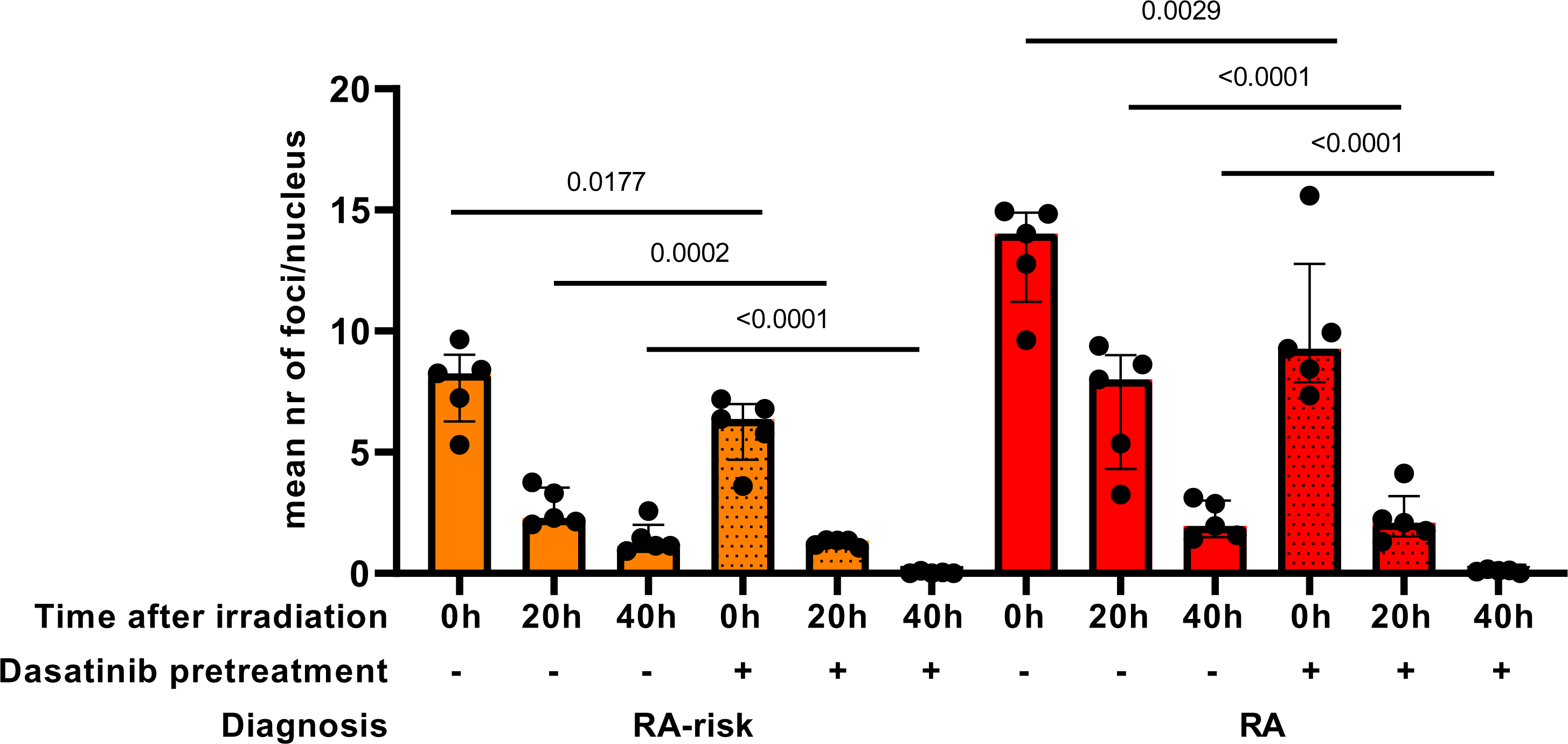
Dasatinib treatment significantly improves irradiation induced DNA damage repair in cultured LNSCs. A) Representative images of yH2AX foci (red) and DAPI staining (blue) in cultured LNSCs. B) Representative images of yH2AX foci (red) and DAPI staining (blue) in cultured LNSCs after DNA damage induction using gamma-irradiation. C) Average number of yH2AX foci per nucleus in cultured LNSCs directly, 20 hours and 40 hours after irradiation. Mean value per donor was determined through quantification of Z-stack images of approximately 50 cells per donor. All donors passage 7, N=5 per group. Data are presented as median + interquartile range. Statistical differences were determined using 2-way ANOVA + Dunnett’s T3 multiple comparisons test.

## ABBREVIATIONS

RA: rheumatoid arthritis
RF: rheumatoid factor
ACPA: anti-citrullinated protein antibodies
RA-risk: individuals who are positive for RF or/and ACPA and at risk of developing RA
LNSC: Lymph node stromal cell

## Acknowledgements

We thank our study subjects for participating in the study, the AMC radiology department for lymph node tissue sampling, the rheumatology lab for sample processing and Tatum van Maanen for the DNA damage analysis support.

## Contributors

TJ and LB designed the study. LB, JS, JB, CG, PK and TJ substantially contributed to the data acquisition. LB, TJ, RH and PK contributed to the statistical analysis and interpretation of the data. LB and TJ wrote the manuscript. All authors critically reviewed the manuscript and approved the final version for submission. LB acts as guarantor for this work.

## Patient and public involvement statement

This research was done without formal patient and public involvement. However, obtained results are shared with study participants during regular patient information meetings where study participants and others can ask questions and discuss our findings with researchers involved.

## Patient consent for publication

Not applicable

## Ethical approval

The study was performed according to the principles of the Declaration of Helsinki, approved by the Institutional Review Board of the Academic Medical Center and all study subjects gave their written informed consent.

