## Supplementary methods for "Senescence phenotype of lymph node stromal cells from patients with rheumatoid arthritis is partly restored by dasatinib treatment"

**Study subjects**

Individuals with arthralgia and/or a family history of RA who were positive for anti-citrullinated protein antibodies (ACPAs; detected by the anti–cyclic citrullinated peptide [anti–CCP] antibody test (CCPlus anti-cyclic citrullinated peptide 2 ELISA (ULN 25 kAU/L) Eurodiagnostica, Nijmegen, the Netherlands) and without any evidence of arthritis upon examination were included. These individuals were considered to be at risk of developing RA (RA-risk individuals), characterized by the presence of systemic autoimmunity associated with RA, but without clinical arthritis (defined as phase c+d, according to EULAR recommendations) (1, 2). After median follow-up of 20 months (11-46 (IQR)) none of these individuals had developed RA despite the presence of ACPAs. However, we expect that 30% of these individuals will eventually develop arthritis within 3-4 years (3-5). For comparison, we included patients diagnosed with RA (ACR/EULAR 2010 criteria (2)) who were all biological naïve, ACPA positive and while two patients had longstanding RA (>8 years), three had a recent diagnosis. Finally, aged-matched seronegative healthy volunteers without any history of autoimmunity or inflammatory disease and no present or previous use of disease-modifying anti-rheumatic drugs (DMARDs) or biologicals were included. Study subjects were recruited either via the outpatient clinic of the Department of Rheumatology and Clinical Immunology at the Amsterdam UMC, via referral from the rheumatology outpatient clinic of Reade, Amsterdam, or via testing family members of RA patients in the outpatient clinic. LN tissues were collected by ultrasound-guided inguinal LN needle core biopsy as previously described (6). The study was performed according to the principles of the Declaration of Helsinki (7), approved by the Institutional Review Board of the Amsterdam UMC and all study subjects gave their written informed consent.

**Flow cytometry for direct ex vivo stromal cell analyses**

For direct *ex vivo* comparison, LNs from kidney transplantation recipients were collected and carefully cleaned of fat and connective tissue. Subsequently, LNs were cut into small pieces (<0.5cm^2^) after which LN tissue samples (from kidney transplantation and inguinal core needle biopsies) were enzymatically digested for 3 times 15 minutes. Digestion mix consisted of 0.2mg/mL collagenase P, 0.8mg/mL dispase II and 0.1mg/mL DNase I (all Roche, Woerden, The Netherlands), in RPMI (Invitrogen, Landsmeer, The Netherlands). After every digestion step, the cell suspension was filtered through a 100µm nylon cell strainer (BD Falcon, San Jose, CA) and after the final step cells were washed and collected in PBA buffer (PBS containing 0.5% BSA and 0.01% NaN_3_ (Sigma Aldrich, Zwijndrecht, The Netherlands). Next, cells were stained for 60 minutes at 4°C with unconjugated anti-PDPN (NC-08, Angiobio). After incubation, cells were washed twice in PBA buffer and stained for 30 minutes with fluorochrome-labeled antibodies against the following markers: CD31-Alexafluor488 (clone WM-59, Biolegend, San Diego, CA), HLA-DR-PE (clone L243, eBioscience), Alexa Fluor 647 (polyclonal, Invitrogen), CD235a-eFluor450 (clone HIR2, eBioscience), CD45-eFluor450 (clone HI30, eBioscience), PD-L1-BV510 (clone 29E.2A3), CD86-BV650 (clone IT2.2, Biolegend), CD80-PEcy7 (clone 2D10, Biolegend), CD40-PEdazzle594 (clone 5C3, Biolegend), CD11c-AlexaFluor700 (clone Bu15, Biolegend), Viability dye-eFluor780 (Invitrogen). Subsequently, cells were washed twice in PBA buffer and analyzed using the BD FACS Aria SORP Cell Sorter. Flow cytometry data was analyzed using FlowJo 10.8.1 (Tree Star, Ashland, OR).

**Stromal cell culture, senescence induction and dasatinib treatment**

LNSCs were isolated and expanded *in vitro* as previously described resulting in cultures containing lymph node fibroblasts (8). Experiments were performed using cultured human LNSCs between passages 5 and 7. To induce DNA damage, control LNSCs were irradiated in culture flasks at 10Gy using Cellrad+ (Precision X-ray, Madison, CT) and passaged simultaneously with non-irradiated cells from the same donor. RA-risk, RA and irradiated control LNSC were treated with 5µM dasatinib (MedChemExpress, Monmouth Junction, NJ) for 24h to remove senescent cells. After 24h treatment, culture flasks were washed with complete cell culture media and cultured until 80% confluence based on the flask containing untreated cells after which all cells were simultaneously harvested for analyses.

For analyses, cultured LNSC were non-enzymatically detached with TrypLE Select (Gibco, Bleiswijk, The Netherlands) for 7 minutes at 37°C and collected for flow cytometry experiments. Cells were washed and collected in PBA buffer (PBS containing 0.5% BSA and 0.01% NaN_3_ (Sigma Aldrich, Zwijndrecht, The Netherlands). Subsequently, LNSCs were measured on a Spectral Analyzer SP6800 (Sony Biotechnology, Weybridge, United Kingdom) and analyzed using FlowJo 10.8.1 (TreeStar Inc., Ashland, OR). Cell size and granularity were determined based on the parameters FSC-W and SSC-A respectively.

**Quantitative real-time PCR**

Total RNA was isolated using the RNA micro kit (Qiagen, Venlo, The Netherlands) according to manufacturer’s instructions. Subsequently, cDNA was prepared using the RevertAid H Minus First Strand cDNA Synthesis kit (Thermo Fisher Scientific, Landsmeer, The Netherlands). Quantitative PCR was performed using either Taqman® Gene Expression master mix combined with Taqman assays or fast SYBR® Green PCR master mix (all Applied Biosystems, Life Technologies, Zwijndrecht, The Netherlands) combined with in house designed primers (Thermo Fisher). Primer sequences and Taqman assays are described in table 2. For detection we used QuantStudio 3 (Applied Biosystems). Values of each target gene were normalized by the geometric mean expression levels of two reference genes; *RPLP0* and *POLR2G*. An arbitrary calibrator sample was used to correct for inter-plate differences. To calculate the relative quantity (RQ), the standard curve method was applied to SYBR green assays while the delta-delta Ct method was applied to Taqman assays.

| **Table 2. Overview of primer sequences** | | | | |
| --- | --- | --- | --- | --- |
| **SYBRgreen** | | | | |
| **Gene** | **mRNA transcriptID** | | **Forward sequence** | **Reverse sequence** |
| *POLR2G* | | NM_002696.3 | GAGGTCGTGGATGCTGTTGT | TCTCTGAAGGGATGGAATGTCG |
| *RPLP0* | | NM_001002.4 | GCAGCATCTACAACCCTGAAGT | GCAGACAGACACTGGCAACAT |
| *FOXO4* | | NM_001170931.1 | GGAAAAGGCCATTGAAAGCG | ATGAACTTGCTGTGCAGGGA |
| *TP53* | | NM_001126116.1 | CAGTCACAGCACATGACGGA | GCCAGACCATCGCTATCTGAG |
| *CDKN1A* | | NM_078467.2 | AGACCAGCATGACAGATTTCTACC | GCGGATTAGGGCTTCCTCTT |
| *CDKN2A* | | NM_000077.4 | TCCCTCAGACATCCCCGATT | CCTGTAGGACCTTCGGTGAC |
| *GLB1* | | NM_001079811.2 | TTTGCTCTGCGAAACATCATCC | GCTCCCACTGTCTTTAACTTTTCC |
| *BCL2L1* | | NM_138578.3 | CTGTGCGTGGAAAGCGTAGA | GCTGCTGCATTGTTCCCATAG |
| *SIRT1* | | NM_001314049.1 | GAGCAGATTAGTAGGCGGCTT | CTCAGCGCCATGGAAAATGT |
| *NAMPT* | | NM_005746.3 | TCTGGAAACCCTCTTGACACTG | GTTTCATGCCTTCTACAATCTCTTG |
| *PARP1* | | NM_001618.4 | AACCGAAGATTGCTGTGGCA | ACCAAACATGTAGCCTGTCACG |
| *CD38* | | NM_001775.4 | GGTGGAAGAGAAGATTCCAGAGAC | TAAAACAACCACAGCGACTGG |
| *EFNB1* | | NM_004429.5 | GGAGGCAGACAACACTGTCA | TCCTGGTTCACAGTCTCATGC |
| *LMNB1* | | NM_001198557.1 | AAATTCTCAGGGAGAGGAGGT | TTGGATGCTCTTGGGGTTC |
| **Taqman assays** | | | | |
| **Gene** | | **Assay ID** |  |  |
| *POLR2G* | | Hs00275738_m1 |  |  |
| *RPLP0* | | Hs00420895_gH |  |  |
| *NOTCH3* | | Hs01128537_m1 |  |  |

**Senescence, DNA damage and repair analyses**

LNSCs were seeded in triplicate at a density of 25’000 cells/well on sterilized round 12mm coverslips (Knittel-Glaeser, Bielefeld, Germany) in 24-wells plates (Corning, Amsterdam, The Netherlands) and cultured for 24h in complete culture medium. After 24h, cells were washed with PBS and fixed using 4% paraformaldehyde (PFA) for 10 minutes. After washing with PBS, cells were permeabilized for 60 minutes at room temperature (RT) with PBS containing 1% BSA and 0.1% saponin (Sigma-Aldrich) and stained overnight at 4°C with monoclonal mouse IgG anti-human yH2AX (Ser139, Sigma-Aldrich). Next day, cells were washed with PBS + 1% BSA + 0.1% saponin and incubated for 30 minutes at 4°C with secondary anti-mouse IgG1 AlexaFluor633 (Invitrogen, Landsmeer, the Netherlands). Subsequently, coverslips were removed from the wells and left to dry at RT in the dark. Coverslips were mounted to microscope slides with DAPI Vectashield Hardset mounting media (Vector Laboratories). yH2AX staining representing DNA damage at baseline was analyzed using confocal microscopy (TCS SP8, Leica Microsystems). DNA damage repair analysis was performed on irradiated cells (1Gy) followed by yH2AX staining either directly, 3h or 24h after irradiation. DNA damage repair was visualized using a Leica DMi8 microscope and staining intensity was analyzed in approximately 50 cells/donor using LAS X 3D and QuPath (v0.3.2).

Lysosomal content was measured by staining for lipofuscin granules and senescence associated β-galactosidase (SA-β-gal) activity in cultured LNSC. Lipofuscin granules were stained using SenTraGor™ reagent (Lab Supplies Scientific, Athens, Greece). 25’000 LNSC/well (for lipofuscin) or 10’000 LNSCs/well (for SA-β-gal) were seeded in triplicate on round 12mm coverslips in 24-wells plates and cultured in a 37°C/5% CO_2_ incubator for 24h. After 24h, immunofluorescent staining of SenTraGor™ was performed according to the manufacturer’s instructions (9). SA-β-gal was stained using the senescence detection kit (Abcam, Cambridge, UK) according to the manufacturer’s instructions. Subsequently, coverslips were removed from the wells and mounted to microscope slides with DAPI Vectashield Hardset mounting media. For analysis of lipofuscin granules and SA-β-gal approximately 50 cells/donor were imaged using confocal (TCS SP8, Leica Microsystems, Amsterdam, The Netherlands) and wide-field microscopy (DM6, Leica) and analyzed using Las X 3D and the Senescence Counter macro in ImageJ (10).

**Cellular viability and live cell analysis of proliferation and migration**

Cellular viability was measured using MTT reagent (3-[4,5-Dimethylthiazol-2-yl]-2,5-diphenyltetrazolium bromide; Thiazolyl blue, Sigma)). LNSC were seeded at 2’500 cells/well in 96-wells plates and an MTT assay was performed 24, 72 and 120 hours after seeding. MTT stock solution (5mg/mL) was added in 1:10 dilution of the original culture media volume and incubated for 2 hours. After incubation, culture medium was removed and converted dye was solubilized using acidic isopropanol, 4mM HCl + 0.1% NP-40 (Calbiochem, San Diego, CA) in absolute isopropanol. Absorbance of converted dye was measured using a spectrophotometer (VersaMax, Molecular Devices, San Jose, CA) at 580nm.

Cellular proliferation and migration of cultured LNSCs was followed over time using the IncuCyte® S3 ZOOM live cell imaging microscope. For proliferation, cells were seeded at a density of 2’500 cells/well in 96-wells plates and placed in the IncuCyte® for 6 days. For migration, LNSCs were seeded at 10’000 cells/well in a 96-well ImageLock plate (Sartorius, Goettingen, Germany) and cultured overnight in a 37°C/5% CO_2_ incubator. The next day, a scratch wound was created using the IncuCyte® wound maker according to the manufacturer’s instructions. After wounding, cells were washed with PBS and fresh complete cell culture media was added. Culture plates were placed in the IncuCyte® for 5 days. Relative wound density, reflecting the ratio of the occupied area to the total area of the initial scratched region, and confluence were analyzed using the Incucyte software.

**Statistics**

Data are presented as median with interquartile range (IQR). Statistical differences between the study groups at baseline were analyzed using a Kruskal–Wallis test followed by a post-hoc Dunn’s test, a two-way ANOVA or repeated measures ANOVA test with Geisser-Greenhouse correction followed by a Dunnett’s multiple comparison test, where appropriate. A Wilcoxon matched pairs signed rank test was used to analyze the effect of irradiation and dasatinib treatment. GraphPad Prism software (version 9.1.0, La Jolla, CA, USA) was used for statistical analysis. p-values < 0.05 were considered statistically significant.

**Supplementary figure 1. Selecting potential senolytics for LNSCs.** A) Viability of RA and control LNSCs after navitoclax treatment for 24 hours. B) Relative expression level of *BCL2L1* mRNA. A gene related to the anti-apoptotic pathway of BCL-XL and a target of navitoclax. C) Viability of LNSCs after piperlongumine treatment for 24h. D) Viability of LNSCs after quercetin treatment for 24h. E) Viability of LNSCs after dasatinib treatment for 24h. Viability was measured using MTT assays. F) Relative expression level of *EFNB1* mRNA, a gene encoding for ephrin B1 which is a type I membrane protein and ligand for dasatinib. All donors passage 6 for qPCR analysis, N=5 per group. Data are presented as median + interquartile range. Statistical differences were determined using a Kruskal-Wallis followed by Dunn’s multiple comparisons test.

**Supplementary figure 2. Gating strategy and flow cytometry plots of representative donors.** A) Flow cytometry gating strategy used to identify single cells and LNSCs. Numbers adjacent to the outlined areas indicate percentages of cells in the gated population. B) Representative flow cytometry plots of irradiated and dasatinib (das) treated LNSCs. C) Representative mean fluorescent intensity (MFI) plots of FITC autofluorescence in unstained LNSCs. Controls (green), RA-risk individuals (orange) and RA patients (red).

**Supplementary figure 3. Gene expression profiles in LNSCs.** Gene expression levels of senescence associated genes in cultured LNSCs. All donors passage 6, N=5 per group. Data are presented as median + interquartile range. Statistical differences at baseline were determined using a Kruskal-Wallis test followed by Dunn’s multiple comparisons test and a Wilcoxon matched pairs signed rank test was used to analyze the effect of dasatinib treatment.

**Supplementary figure 4. Dasatinib treatment significantly reduces cell viability in RA-risk LNSCs.** A) Viability of irradiated and dasatinib treated LNSCs measured using MTT. B) Real-time cellular proliferation of irradiated and dasatinib treated LNSCs measured using IncuCyte software. C) Real-time cellular migration capacity of irradiated and dasatinib treated LNSCs measured using scratch wound assay software from IncuCyte. All donors passage 6, N=5 per group, 3 technical replicates per condition for viability and 5 technical replicates for proliferation and migration assays. Data are presented as median + interquartile range. Statistical differences were determined using a repeated measures ANOVA with the Geisser-Greenhouse correction followed by Dunnett’s multiple comparisons test.

**Supplementary figure 5. Dasatinib treatment significantly improves irradiation induced DNA damage repair in cultured LNSCs.** A) Representative images of yH2AX foci (red) and DAPI staining (blue) in cultured LNSCs. B) Representative images of yH2AX foci (red) and DAPI staining (blue) in cultured LNSCs after DNA damage induction using gamma-irradiation. C) Average number of yH2AX foci per nucleus in cultured LNSCs directly, 20 hours and 40 hours after irradiation. Mean value per donor was determined through quantification of Z-stack images of approximately 50 cells per donor. All donors passage 7, N=5 per group. Data are presented as median + interquartile range. Statistical differences were determined using 2-way ANOVA + Dunnett’s T3 multiple comparisons test.
